# A novel ionic model for matured and paced atrial–like hiPSC–CMs integrating *I*_*Kur*_ and *I*_*KCa*_ currents

**DOI:** 10.1101/2024.01.12.574782

**Authors:** Sofia Botti, Chiara Bartolucci, Claudia Altomare, Michelangelo Paci, Lucio Barile, Rolf Krause, Luca Franco Pavarino, Stefano Severi

## Abstract

Human induced pluripotent stem cells–derived cardiomyocytes have revolutionized the field of regenerative medicine, offering unparalleled potential for *in–vitro* modeling of normal and pathological human cardiomyocytes. The ability to produce stem cardiac myocytes in abundance has opened new avenues for drug efficacy and safety testing, as well as the study of conditions such as atrial fibrillation, a familial cardiac disorder. The development of atrial fibrillation is influenced by ion channel mutations, genetic variants, and other risk factors. Stem cells derived cardiomyocytes hold promise in personalized medicine, as they share the genetic heritage of the donor. While mathematical models have focused on immature stem cardiomyocytes phenotypes, they have primarily relied on a system of stiff ordinary differential equations. Computational modeling of diseased tissue presents an opportunity to evaluate drugs in a patient-specific manner, thereby improving therapeutic targets and ablation techniques. Previous studies categorized cell phenotypes based on action potential morphology, yet classification criteria remains ambiguous.

This work introduces the first atrial-specific *in–silico* model of stem cells ionic currents, leveraging experimental data provided by Altomare et al. It begins by summarizing the baseline electrophysiological model and mathematical descriptions of atrial–specific additional currents. Model parameter tuning was performed through automatic optimization techniques to ensure realistic action potential shape and expedite the parameter adjustment process. The resulting model was validated against rate dependence and atrial–specific ion current blocking data. In summary, the development of an atrial-specific *in–silico* model represents a significant step forward in understanding cardiac electrophysiology and the potential for personalized medicine in treating conditions like atrial fibrillation. This model offers new tools for drug evaluation, therapeutic improvement, and a deeper comprehension of cardiac phenotypes.

**Author summary:** Human induced pluripotent stem cells have revolutionized regenerative medicine since their discovery in 2006, leading to a Nobel Prize in 2012. This kind of pluripotent cells can give rise to different types of specific tissue cells, such as derived cardiomyocytes. Differentiated cardiac cells offer an unlimited supply for studying human heart cells in normal and disease conditions, aiding a patient–specific drug testing and helping to explore pathogenic mechanisms behind different cardiomyopathies, including atrial fibrillation. Atrial fibrillation is a common heart condition, and stem cells with the same genetic heritage as the donor, are ideal for patient-specific treatments.

Recent advances have produced mathematical models for the ionic currents in cardiomyocytes derived from stem cells, focusing on immature forms and enabling virtual drug testing. However, previous models did not capture the atrial–specific characteristics. We decided to create and introduce by this study the first atrial–like *in–silico* model for these cells, using novel experimental data. Thus, we describe the baseline model and additional atrial–specific currents, we tune the model parameters using automatic optimization technique, and we validate the model’s accuracy in simulating atrial action potentials and ion current blockage. This research paves the way for better understanding and treating atrial fibrillation and other heart conditions.

## Introduction

Human induced pluripotent stem cells–derived cardiomyocytes (hiPSCs) have enormously advanced the field of Regenerative Medicine since their discovery in 2006 by Yamanaka et al., [1, 2], which then led to the Nobel Prize in Medicine in 2012. In the same decade, Yamanaka group refined the capacity to differentiate hiPSCs into disease–relevant cell types such as cardiomyocytes (hiPSC–CMs), providing an unprecedented opportunity for the generation of human patient–specific cells for use in disease modeling, personalized drug screening, and regenerative approaches toward precision medicine. The unlimited production of hiPSC-CMs provides new opportunities to evaluate *in–vitro* models of human cardiomyocytes in normal or pathological conditions that can be used in drug efficacy and safety testing. Moreover, hiPSC–CMs also have the potential to become an essential tool to better understand the familial form of atrial fibrillation (AF), a common disease affecting atrial cells, see [3, 4]. The best option currently used for treating the disease is interventional therapy, namely ablation. Several ion channel mutations, along with a range of other genetic variants and broader risk factors, are known to increase the likelihood of developing AF. Therefore, hiPSC-CMs technology perfectly fits the patient–specific medicine challenge, since these cells have the same genetic heritage as the donor.

In recent years, mathematical models of the hiPSC–CMs ionic currents have focused on immature phenotypes, developing a system of stiff ordinary differential equations (ODEs). Previous studies characterizations of the CMs phenotype were based on action potential (AP) morphology, but the classification criteria were still undefined. Thus, the forerunner Paci2013 [5] model was based on recordings obtained from a mixture of ventricular–like (VL), atrial–like (AL), and nodal–like hiPSC-CMs and the phenotypical heterogeneity was reproduced considering different scalings, instead of phenotype–specific currents.

Nowadays, the employment of maturation techniques highlight the chamber-specific AP phenotype of the cells ([6, 7]). Between developed chamber–specific cultures, atrial ones accurately reflect the electrophysiological characteristics of atrial tissue, prove essential in drug testing scenarios. They enable a more nuanced evaluation of drug responses, especially in the context of AF, atrial–specific arrhythmias or pathologies, providing a valuable platform for assessing the safety and efficacy of pharmaceutical interventions in a chamber–specific manner. Furthermore, computational models of a diseased tissue could be used to virtually evaluate drugs through a patient-specific model, by paving the way for improving therapeutic targets or ablation techniques. To this end, a phenotype–specific *in–silico* models, for both AL and VL hiPSC–CMs could be useful to realize a virtual platform, which allows to predict the drug effect in a single cell or in cardiac tissue.

In this direction, the latest version Paci2020 [8] still provides a developed VL model, while in this work, we present the first AL *in–silico* model of the hiPSC–CM ionic currents, based on experimental data provided by [9]. First of all, the baseline electrophysiological model is summarized, as well as the mathematical description of atrial–specific additional currents. Moreover, the fine tuning of the model parameters was performed by means of an automatic optimization technique, in order to reproduce realistic AP transient shape and to speed up the parameter tuning phase. Finally, the resulting model was presented and validated against rate dependence and atrial–specific ion current blocking data.

## Materials and methods

### Experimental data set

In cultures of hiPSC–CMs, the time independent inward-rectifier K^+^ current (*I*_*K*1_), that usually maintains negative the membrane diastolic potential (MDP), current can be too low or even lacking, leading to unstable MDP, if compared to the mature CMs. These immature electrophysiological conditions correspond to a spontaneous firing activity or a depolarized resting (*≃-*20 mV). Dynamic clamp (DC) is a valid and effective approach to overcome immature electrical properties of hiPSC–CM through the injection of a virtual *I*_*K*1_ current in a real time mode. DC then leads to a more hyperpolarized derived–CMs MDP allowing the cells to exhibit a more mature AP.

The experimental data set consists of AP recordings from hiPSC–CMs obtained by whole cell patch clamp configuration, in paced condition, at room temperature (37^*o*^C) and with extracellular concentrations Na^+^ = 154.0, K^+^ = 4.0, Ca^2+^ = 2.0 mM. The hiPSCs were differentiated into cardiomyocytes and treated with retinoic acid (RA, 1 μM) to induce atrial differentiation, [7, 9–11]. During the experiments, APs, recorded from the hiPSC–CMs, were acquired to drive the numerical formulation of the time independent *I*_*K*1_ current (taken from the Koivumäki atrial AP model [12]) in DC. Modelled *I*_*K*1_ was calculated in real–time and injected into the myocyte during continued AP recording. All experimental APs data were corrected for the estimated liquid junction potential (a shift of *-*8 mV was applied), see [13].

The following biomarkers, summarized in Table 1, were considered: MDP, action potential amplitude (APA), AP duration (APD) at 20, 50, and 90% of AP repolarization (APD_20_, APD_50_, and APD_90_), maximum upstroke velocity (V_max_) and APD_20*/*90_ (APD_20_ over APD_90_) ratio selected as the critical biomarker to discriminate AL *versus* VL hiPSC–CM. Each measurement was characterized by its mean value and its standard deviation (Std. Dev.) over a variable number of beats on a total of 10 cells.

**Table 1.**
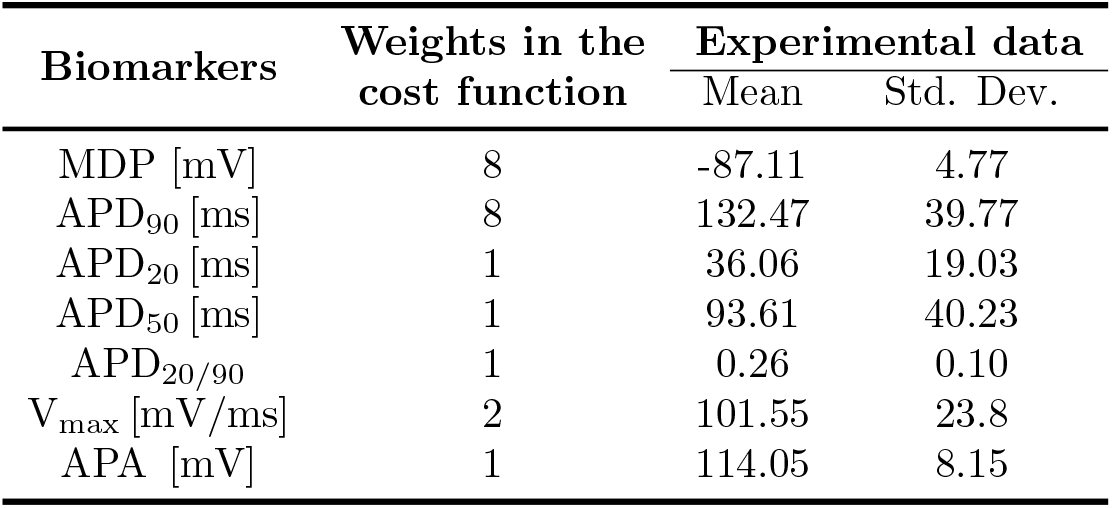
Experimental data. Experimental data at 1 Hz provided by Altomare et al. in [9]. These biomarkers were also considered for the parameter optimization. Weights used in the automatic optimization process are also summarized in the table.

| Biomarkers | Weights in the cost function | Experimental data |  |
| --- | --- | --- | --- |
|  |  | Mean | Std. Dev. |
| MDP [mV] | 8 | -87.11 | 4.77 |
| APD<sub>90</sub> [ms] | 8 | 132.47 | 39.77 |
| APD<sub>20</sub> [ms] | 1 | 36.06 | 19.03 |
| APD<sub>50</sub> [ms] | 1 | 93.61 | 40.23 |
| APD<sub>20/90</sub> | 1 | 0.26 | 0.10 |
| V<sub>max</sub> [mV/ms] | 2 | 101.55 | 23.8 |
| APA [mV] | 1 | 114.05 | 8.15 |

### Atrial parametrization of the Paci2020 hiPSC–CM model

The Paci2020 model [8], adapted to the extracellular K^+^ concentration of the experimental recordings of Altomare et al. [9], was used as the baseline to build an atrial specific *in–silico* model for AL hiPSC–CMs. The matching with the experimental condition K_o_ = 4.0 mM affected the reversal potential E_*K*_ = *-*96.8 mV and ion currents, such as *I*_*NaK*_ and *I*_*Kr*_, with effects on the AP duration.

Following the classical Hodgkin–Huxley formalism, the ionic current through membrane channels was described by the transmembrane potential *V*_*m*_, the vector of the R gating variables ***w*** = (*w*^1^,…, *w*^R^), where R = 18 in Paci2020, and ionic concentrations 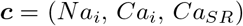, where Ca_*SR*_ means the Ca^2+^ concentration in the Sarcoplasmic Reticulum (SR). This leads to the following system of ODEs.

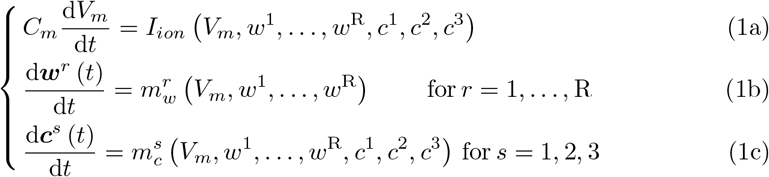

where *C*_*m*_ is the membrane capacitance and *I*_*ion*_ is the sum of 13 membrane currents and the stimulus current.

As a first step towards the atrialization we rescaled 11 parameters of Paci2020 by factors provided by the literature and summarized in Table 1 of Paci2013, [5]. Among them, two different sets can be identified. The first one corresponds to parameters for which a specific relation between VL and AL phenotype is provided in the literature, since they do not derive from data fitting. The corresponding value in Paci2020 is now the ventricular parameter, used to evaluate the atrial one through the proposed scaling in Paci2013. In the second one, parameters were derived by fitting a mixed pool of voltage clamp data, affected by phenotypical heterogeneity. There, the resulting basic fitting parameter was scaled through function *f*_*a*_, *f*_*v*_ to the AL or VL version. In our case, basic fitting parameters needed for the rescaling, are computed through the inverse function 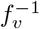 from the VL value provided in Paci2020 [8]. Some of these changes are summarized in Table 2, while parameters not appearing in the table will feed the optimization process, detailed in future sections.

**Table 2.** Atrial reparametrization of Paci2020. First set of parameters, scaled according to the specific relation provided in Paci2013. Setting VL parameters provided in Paci2020 as a starting point, AL values for the presented model can be deduced using the given reparametrization.

|  | Basic fitting | VL parameter | AL parameter | Units |
| --- | --- | --- | --- | --- |
| $C_m$ | $C_{m,v}/1.113$ | $C_{m,v} = 98.7109$ | $0.887 * C_m = 78.6672$ | pF |
| $V_{up}$ | — | $V_{up,v} = 0.82205$ | $0.3924 * V_{up,v} = 0.3226$ | mM/s |
| $g_{irel}$ | — | $g_{irel,v} = 55.808061$ | $1.1109 * g_{irel,v} = 75.4190$ | mM/s |
| $V_c$ | — | $V_{c,v} = 8800$ | $V_{c,a} = 7012$ | $\mu\text{m}^3$ |
| $V_{SR}$ | — | $V_{SR,v} = 583.728$ | $V_{SR,a} = 465.199$ | $\mu\text{m}^3$ |

### Additional atrial–specific currents

The parental AL Paci2013 model was based on mixed recordings, thus any atrial–specific current could be considered into the model. Several membrane currents are only expressed in the atria and we considered the integration in the new AL model of the ultrarapid delayed rectifier current (*I*_*Kur*_) and the small conductance Ca^2+^ activated K^+^ (SK) channel (*I*_*KCa*_).

### Ultrarapid delayed rectifier current formulation

The outward current *I*_*Kur*_ is the first atrial–specific additional current we integrated into the model, since it plays a significant role in human atrial repolarization and is generally characterized by rapidly activation, and slow and partial inactivation. Due to atrial–specific expression, the pharmacological inhibition of *I*_*Kur*_ takes into account the selective atrial APD prolongation with minimal adverse effects in the ventricles.

We considered the Courtemanche’s formulation described in [14] written as follows:

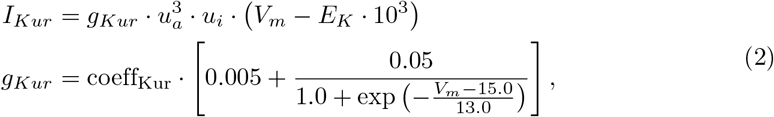

where *g*_*Kur*_ is the maximum conductance and coeff_Kur_ is an additional rescaling factor assumed to be equal to 1. The current dynamic is defined by two specific gating variables, *u*_*a*_, *u*_*i*_, with an Hodgkin-Huxley first order dynamic described by the following equations, where *K*_*Q*,10_ = 3:

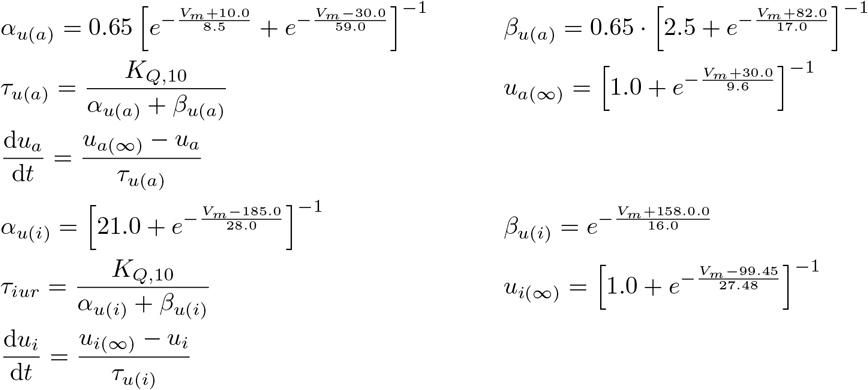

Finally, initial conditions for the new gating variables are given in S1 Appendix.

### Small conductance Ca^2+^ activated K^+^ channel

The second additional current we take into account is *I*_*KCa*_, presented by Skibsbye in 2016, [15]. SK channel opening is described as a two-state Markov model. The opening of the channel simply depends on the sub-sarcolemmal Ca^2+^ concentration, Ca_*i*_. The equations proposed for the opening of the channel are:

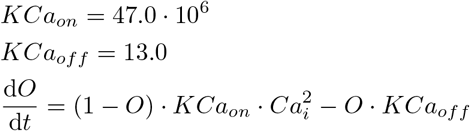

where *O* is the opening gating variable. The resulting current equation is

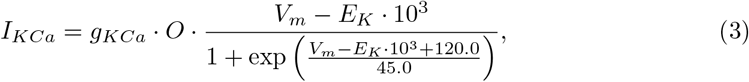

where the maximum conductance is *g*_*KCa*_ = 0.072 [nS/pF].

### Time independent inward–rectifier K^+^ current formulation

As described in the first section, RA–treated CMs were simulates using the DC protocol to overcome the limitation of the low *I*_*K*1_ expression in immature hiPSC-CM, as described in [9, 16, 17]. Since the model has to be consistent with experimental data we are trying to replicate *in–silico* the injection of *I*_*K*1_ current, changing the arbitrarily proposed formulation. Among the two state–of–the–art *I*_*K*1_ *in–silico* currents injected in DC mode [9], we integrated in the model the human atrial specific *I*_*K*1_ formulation published in 2011 by Koivumäki et al. [12], and in 1998 by Nygren et al. [18]. *I*_*K*1_ current, responsible for the late repolarization phase, is thus described by the following equation:

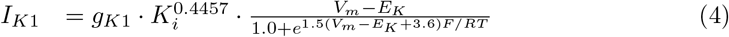

where *g*_*K*1_ = 0.0765 [nS/pF], *F* is the Faraday’s constant, *R* is the gas constant and *T* is the absolute temperature.

### Parameter optimization

The resulting model, extended with additional atrial–specific currents was used as a baseline model. Firstly, we manually increased and calibrated: *(i)* the maximal conductance *g*_*K*1_ in the novel *I*_*K*1_ formulation, mostly responsible of the shortening of the APD_90_ and the hyperpolarization of the MDP, *(ii)* K^+^ driven currents *I*_*Kr*_ and *I*_*Kur*_ affecting the fast repolarization phase, through the maximal conductance *g*_*Kr*_, and the additional scaling factor coeff_Kur_ in equation (2), *(iii)* the maximal conductance *g*_*bNa*_ and the adaptation gate constants of the release *RyR*_*a*1_ in order to restore intracellular Ca^2+^ and Na^+^ concentrations to physiological ranges and prevent the cell from accumulating Ca^2+^ in the SR. Secondly, voltage–dependent inactivation time constants *τ*_*f*1_, *τ*_*f*2_ in *I*_*CaL*_ inactivation gating variable *f*_1_, *f*_2_ where updated in order to shift the recovery from inactivation towards more negative values of *V*_*m*_.

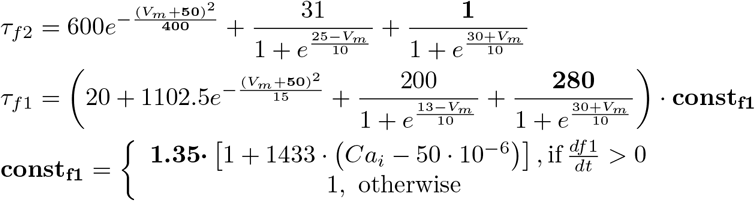

Bold values are resulting optimized parameters.

Finally, we performed the parameter optimization using a hybrid approach, combining a genetic optimization and a simplex optimization, as described in [19]. The first step is usually done using the Matlab function *ga*, based on a genetic algorithm which mimics the natural selection process in biological evolution. The second step corresponds to the Matlab *fminsearchbnd* function, which effectively implements the Nelder–Mead Simplex Method. The resulting hybrid method minimized the cost function, built on the experimental biomarkers we want to simulate. The chosen *in–vitro* biomarkers are the same provided in [9] and they are reported also in Table 1. The cost function structure is defined by the following equations:

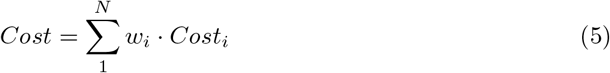

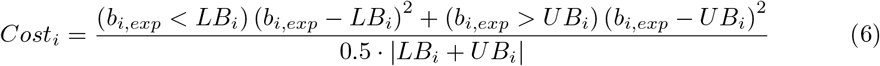

where b_*i,exp*_ is the i^*th*^ biomarker, N the number of biomarkers used, w_*i*_ the weight for each biomarker’s cost and LB_*i*_,UB_*i*_ the lower and the upper bound respectively for the considered biomarker i. Each bound depends on the experimental standard deviation of the single biomarker. Parameters were chosen to include all the main ionic conductances and can be listed as follows: *(i)* the maximum conductances of *I*_*Na*_, *I*_*f*_, *I*_*CaL*_, *I*_*to*_, *I*_*Ks*_, *I*_*pCa*_, *I*_*NaL*_, *I*_*KCa*_, *(ii)* k_*NaCa*_, P_*NaK*_. The parameter values were constrained in a range [*-*30%, +30%] with respect to their starting value, given by Paci2020 model or as a result of its atrial parametrization described in the previous section, in order to avoid non-physiological values, such as negative conductances.

Biomarkers were computed at the steady state (after 800 s) as the average of the last 2 beats. Results are summarized, both for the manual tuning and the automatic optimization, in Table 3.

**Table 3.**
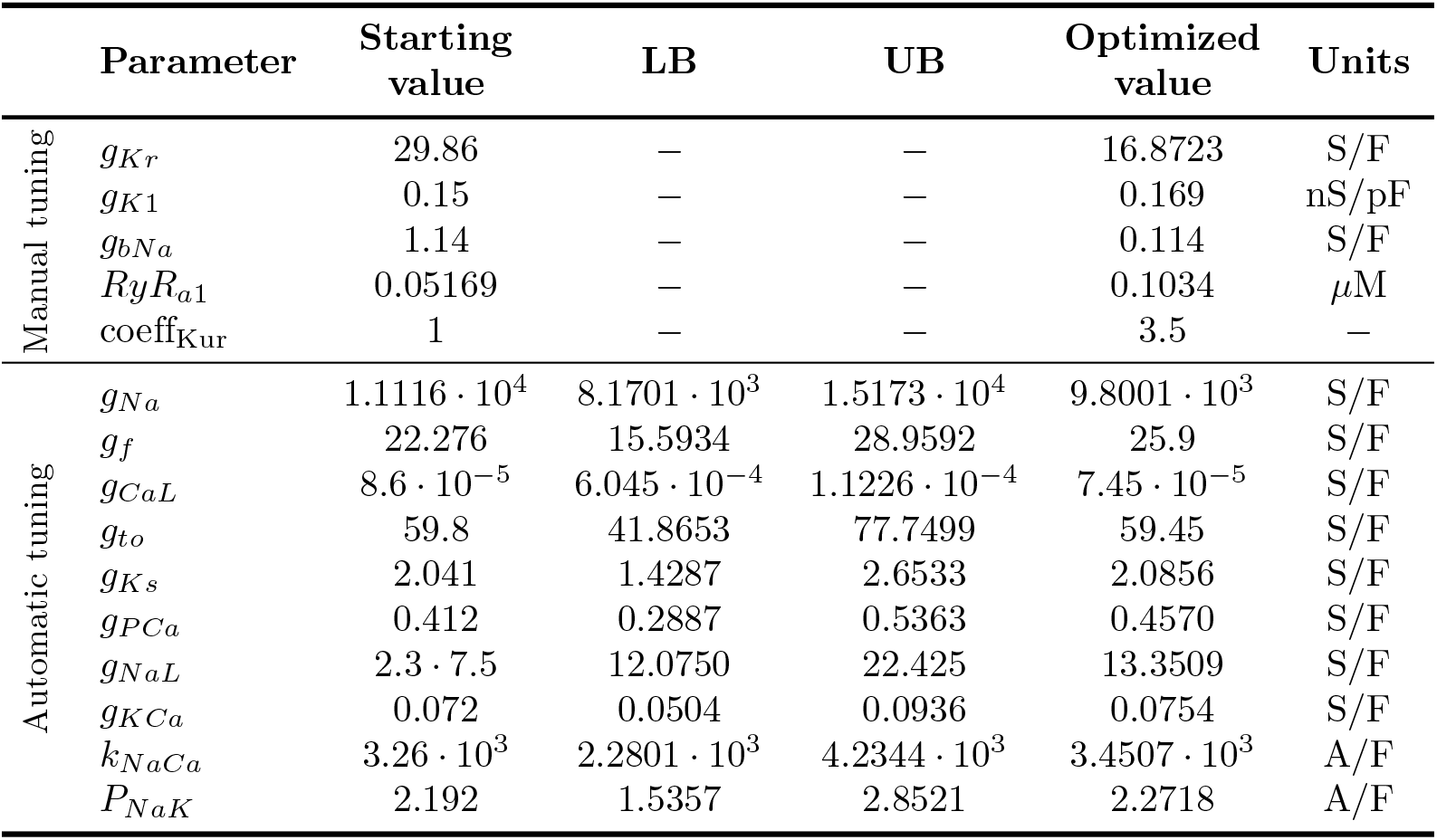
Optimized parameters. We provide the complete set of parameters chosen for manual or automatic optimization. Lower and upper boundaries are provided for the automatic tuning only.

### Numerical simulation setting

Our model was implemented in MATLAB (The MathWorks, Natick, MA). Numerical integration was performed using a solver for stiff systems (ode15s), with an initial step size of 2 *·* 10^−5^ s and a maximum step size of 1 ms. To accurately reproduce experimental condition and the external stimulus applied to the cell, an additional current *I*_*stim*_ (amplitude 1.41 nA, duration 2 ms) was applied in the model with the frequency of 1 Hz. Simulations were always conducted at steady state, reached after 800 seconds of simulations. The resulting initial conditions at steady state are summarized in S1 Appendix. Conversely, to test the APD dependence on the applied pacing rate, we paced the AL model at 1, 1.4, 2 and 4 Hertz, cycle length (CL) 1000, 750, 500 and 250 ms respectively, for 800 beats to reach the steady state. To test the current block effect the model was paced setting the frequency equal to 1 Hz. Model code will be provided under request.

## Results

### Non–mature AL hiPSC–CMs conditions

As discussed in the experimental setup, DC technique allows the cells to reach a stable hyperpolarized MDP in resting conditions and then a physiological AP waveform. During this electronic maturation process, the cell progresses through multiple stages, since the initial unstable depolarized MDP is driven through spontaneous activity and finally reaches the resting.

As a first result, we want that the ionic model in unpaced conditions mimics the maturation process, reaching a stable hyperpolarized MDP. Once achieved the stable resting, is then possible to apply the stimulation protocol and derive the novel AL hiPSC–CMs model. As depicted in Fig. 1, the different stages progession can be perfectly replicated by the unpaced *in–silico* model at different *I*_*K*1_ densities. Indeed, the immature condition, i.e. spontaneous oscillations between depolarized values, corresponds to small percentages of the injected current (*g*_*K*1_ ≤ 0.02 [nS/pF]), while the spontaneous firing activity arises for *g*_*K*1_ ∈ [0.03, 0.06] [nS/pF]. We also note that increasing *g*_*K*1_ values also leads to the increase of the spontaneous firing cycle length. Finally, for values higher than 0.07 [nS/pF] the current leads to an hyperpolarized MDP.

**Fig 1.**
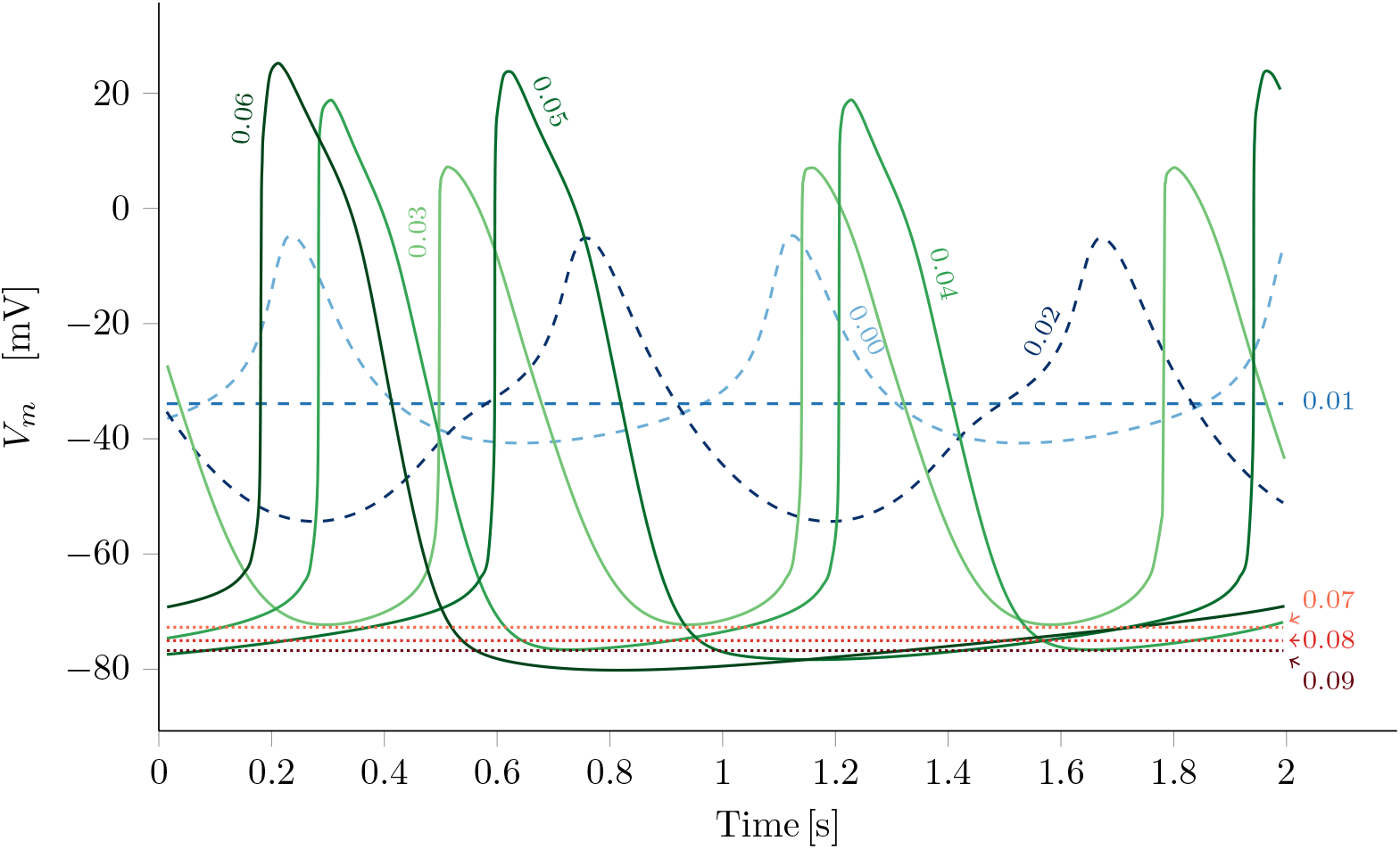
AP morphology with respect to the *g*_*K*1_ variability. The gradual increasing of the *I*_*K*1_ current injection, drives the cell from the initial unstable depolarized MDP (blue dashed lines) to the mature hyperpolarized resting (red dotted lines), passing through the non–mature spontaneous beating (green solid lines). Absolute values of *g*_*K*1_ ([nS/pF]) are displayed near the resulting simulation.

### The new AL hiPSC–CMs model

The introduction of atrial–specific ionic currents and the subsequent automated optimization process successfully identified a new phenotype–specific AL ionic model. A schematic diagram of our AL hiPSC-CM model is reported in Fig 2, showing the cell structure: the model includes two compartments, namely cytosol and SR, as well as the main ion channels, exchangers and pumps. The model follows the classical Hodgkin–Huxley formulation, which describes the transmembrane potential through the ODEs system eq. (1), where R = 20. There, *I*_*ion*_ is the sum of 15 ion currents, exchangers and pumps (see Fig 2), accurately reported in S1 Appendix. All model equations and parameter values are provided in S1 Appendix. In Fig 3 simulated paced AP trace and Ca^2+^ transient at steady state in paced conditions are reported, together with the 15 ionic membrane currents, and the fluxes from the SR.

**Fig 2.**
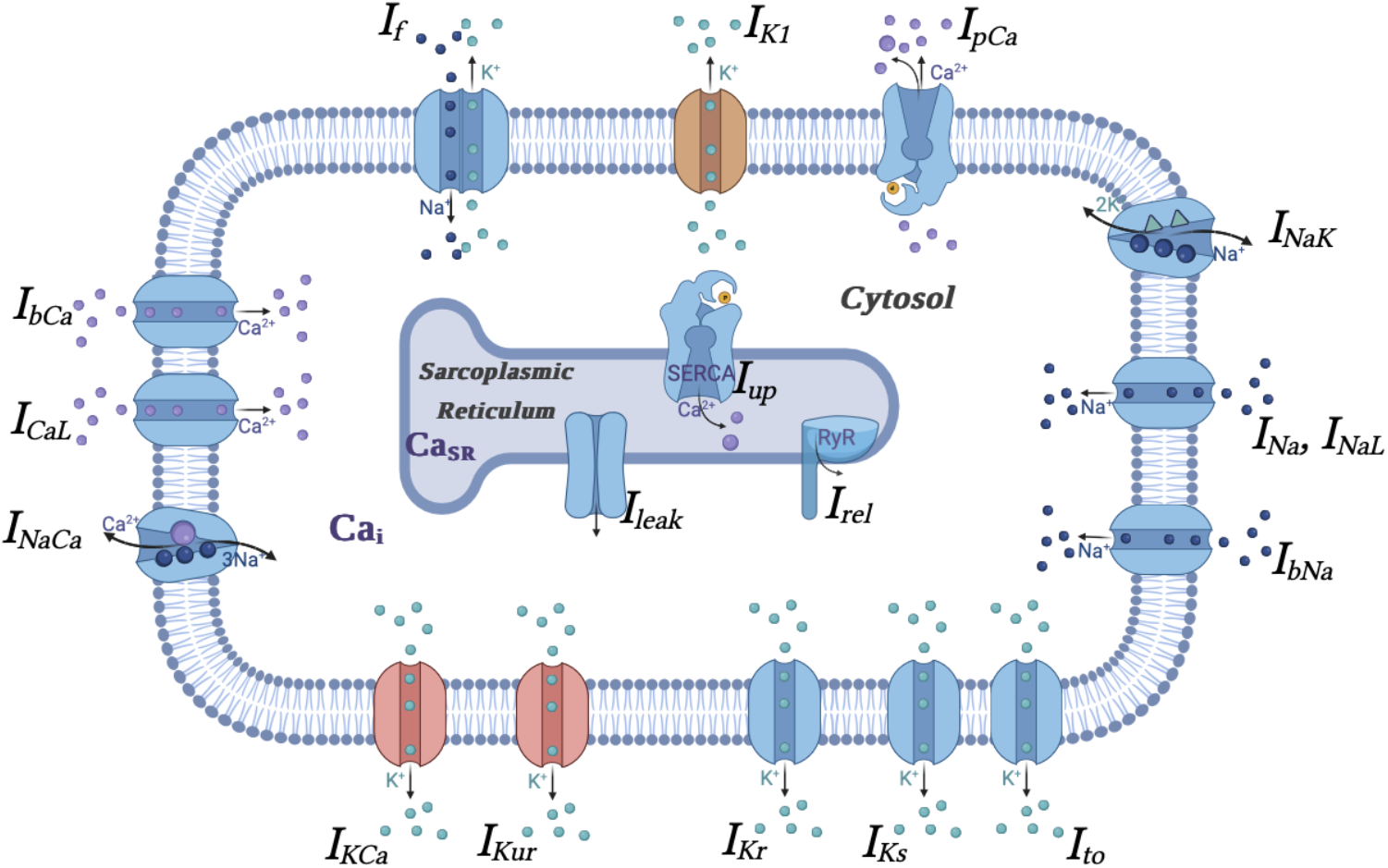
Schematic diagram of the AL hiPSC–CM model. Schematic diagram of the model depicting cell compartments and the major functional components and the 15 membrane currents. Fluid compartments include cytosol and SR. Ca^2+^-handling is described by: *I*_*up*_ uptaking Ca^2+^ by SR, *I*_*rel*_ releasing Ca^2+^ by ryanodine receptors (RyR) and *I*_*leak*_. Pink and brown channels stand for additional atrial–specific currents or with a different formulation, respectively.

**Fig 3.**
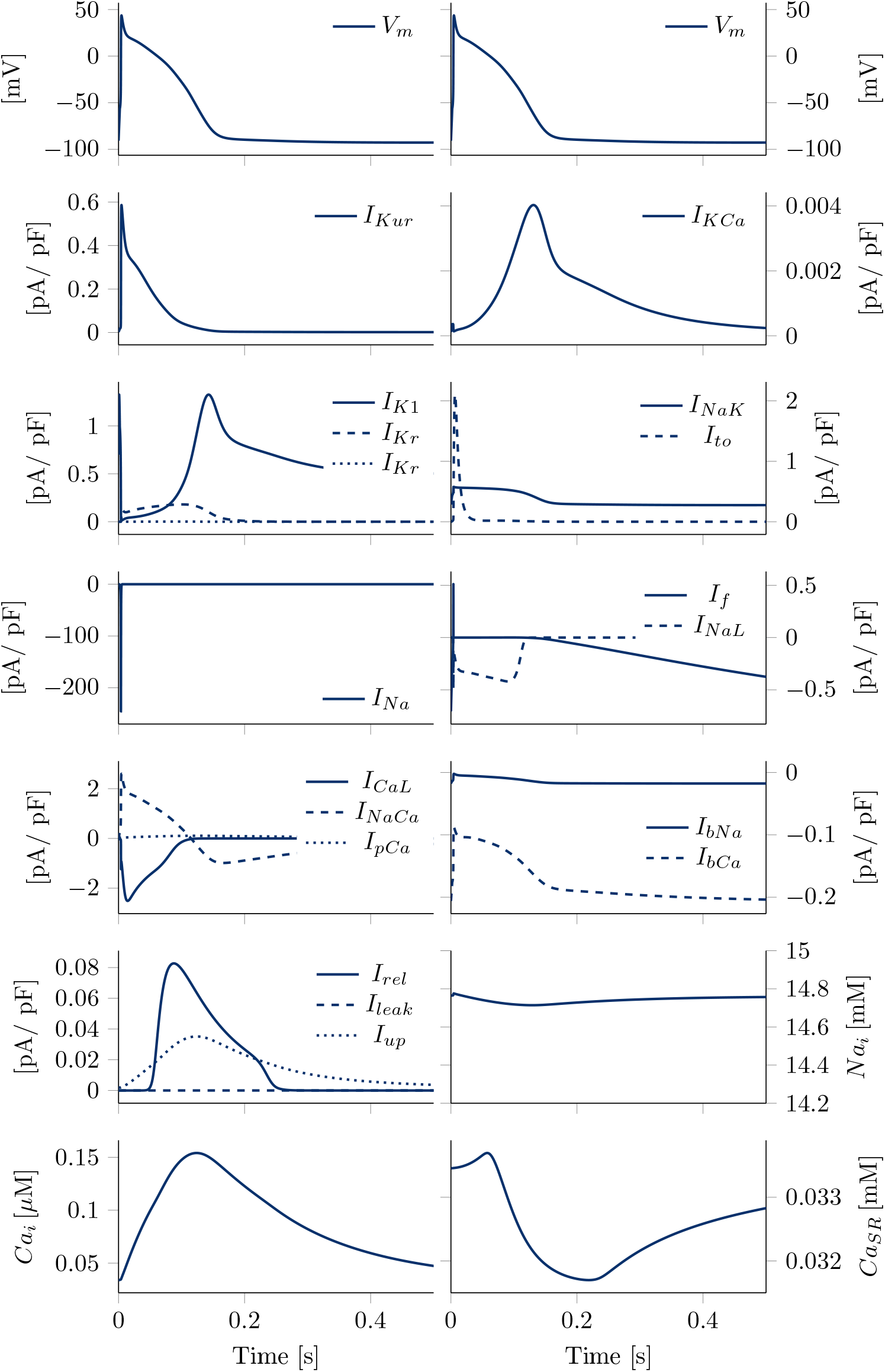
Steady state ionic currents and ionic concentrations dynamic. Ionic currents dynamic, fluxes from the SR, and evolution over time at steady state of the intracellular Na^+^ and Ca^2+^ concentrations and Ca^2+^ concentration in the SR in paced conditions. In the upper panels the transmembrane potential is reported as a reference.

The automated optimization process successfully provides a new set of parameters and identified a new AL model. Fig 4 illustrates the simulated AP and traces from twelve illustrative cells: the comparison highlights that the simulated AP contour is fully in agreement with experimental traces. The simulated AP biomarkers at steady state in paced conditions are in agreement with the *in–vitro* AP biomarker variability ranges, provided as Mean*±*Std. Dev. in [9], as reported in the last column of Table 4, except for the APA, highly dependent on the stimulus current.

**Table 4.** Biomarkers matching. Experimental and simulated values of the biomarkers considered for the parameter optimization. Experimental data are provided as Mean*±*Std. Dev., from [9].

| Biomarkers | Experimental data | Simulated values |
| --- | --- | --- |
| MDP [mV] | $-87.11 \pm 4.77$ | -92.70 |
| APD<sub>90</sub> [ms] | $132.47 \pm 39.77$ | 148.94 |
| APD<sub>20</sub> [ms] | $36.06 \pm 19.03$ | 26.6012 |
| APD<sub>50</sub> [ms] | $93.61 \pm 40.23$ | 96.94 |
| APD<sub>20/90</sub> | $0.26 \pm 0.10$ | 0.1786 |
| V<sub>max</sub> [mV/ms] | $101.55 \pm 23.8$ | 122.503 |
| APA [mV] | $114.05 \pm 8.15$ | 136.53 |

**Fig 4.**
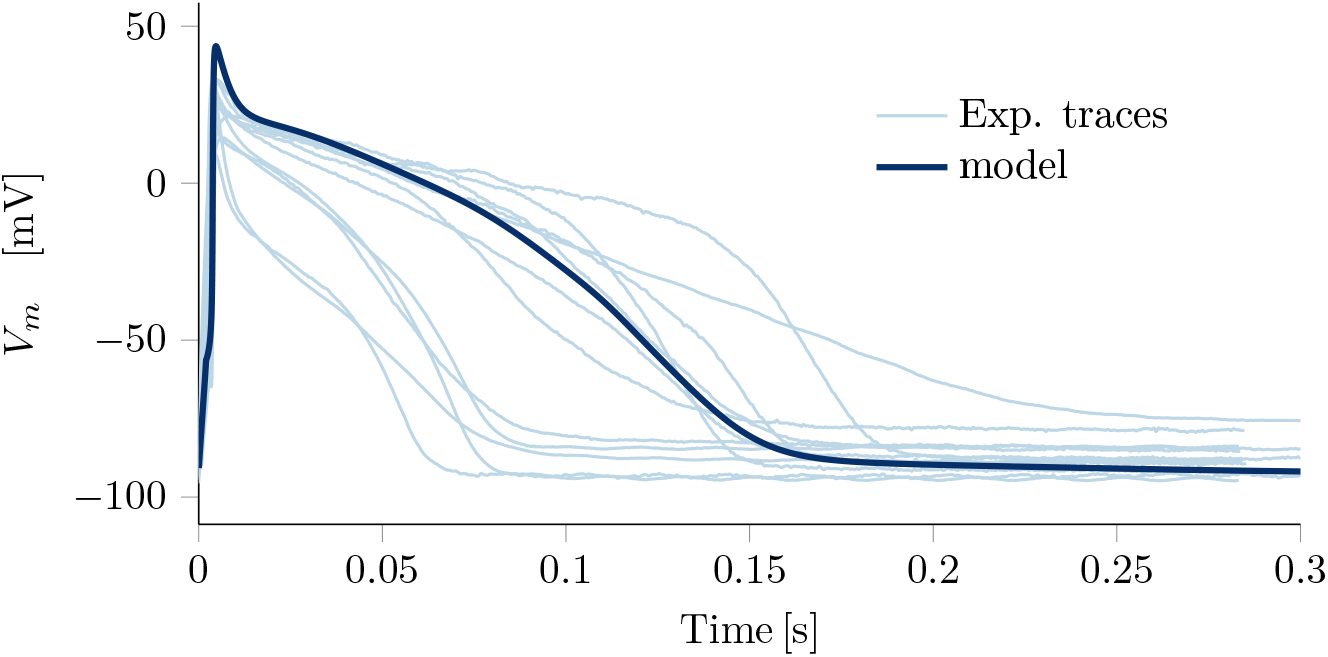
Simulated AP overlapped with experimental traces. Illustrative experimental AP from twelve cells (light blue) and the AP simulated by the AL hiPSC model (dark blue).

### Rate dependence

The dependence of APD on pacing rate is a fundamental property of CMs that, when altered, may promote life–threatening cardiac arrhythmias. In order to validate the new hiPSC–CM model after the introduction of the new atrial–specific currents formulation and the parameters optimization by means of our AP data, we tested the model capability to simulate the APD rate dependence.

To test the APD dependence on the applied pacing rate, the model has been run following the protocol depicted in the previous section, considering different frequencies of stimulation, i. e. CL = 1000, 750, 500, 250. We then compare simulated APD at 90, 50 and 20 % of repolarization with the available *in–vitro* data in [9]. The novel AL model simulations show qualitative agreement with Altomare et al. experiments and traces, as shown in Fig. 5. Furthermore simulated rate dependency curves overlapped with provided experimental error bars depicted in Fig. 6 and a quantitative comparison summarized in Table 5 highlight that APD_90_ and APD_90_ simulated values perfectly fit the experimental ranges. Nevertheless APD_20_ is lower at the highest pacing rates, and this negligible discrepancy is probably due to the APD_20_ dependency on the applied *I*_*stim*_ amplitude.

**Table 5.** Rate dependence. Simulated (Sim.) values with different pacing rates, and experimental (Exp.) data provided as Mean*±*Std. Dev., from [9].

| Biomarkers | CL = 1000 ms |  | CL = 500 ms |  | CL = 250 ms |  |
| --- | --- | --- | --- | --- | --- | --- |
|  | Exp. data | Sim. | Exp. data | Sim. | Exp. data | Sim. |
| APD<sub>90</sub> [ms] | 132.47±39.77 | 148.94 | 109.28±26.60 | 129.8 | 85.25±18.92 | 100.1 |
| APD<sub>20</sub> [ms] | 36.06±19.03 | 26.60 | 29.84±13.79 | 14.85 | 19.30±6.37 | 9.71 |
| APD<sub>50</sub> [ms] | 93.61±40.23 | 96.94 | 75.19±25.67 | 80.85 | 54.36±14.68 | 52.68 |
| APD<sub>20/90</sub> | 0.26±0.10 | 0.18 | 0.27±0.09 | 0.114 | 0.23±0.07 | 0.09 |
| APA [mV] | 114.05±8.15 | 136.5 | 115.8±7.36 | 129.85 | 107.15±7.69 | 132.1 |

**Fig 5.**
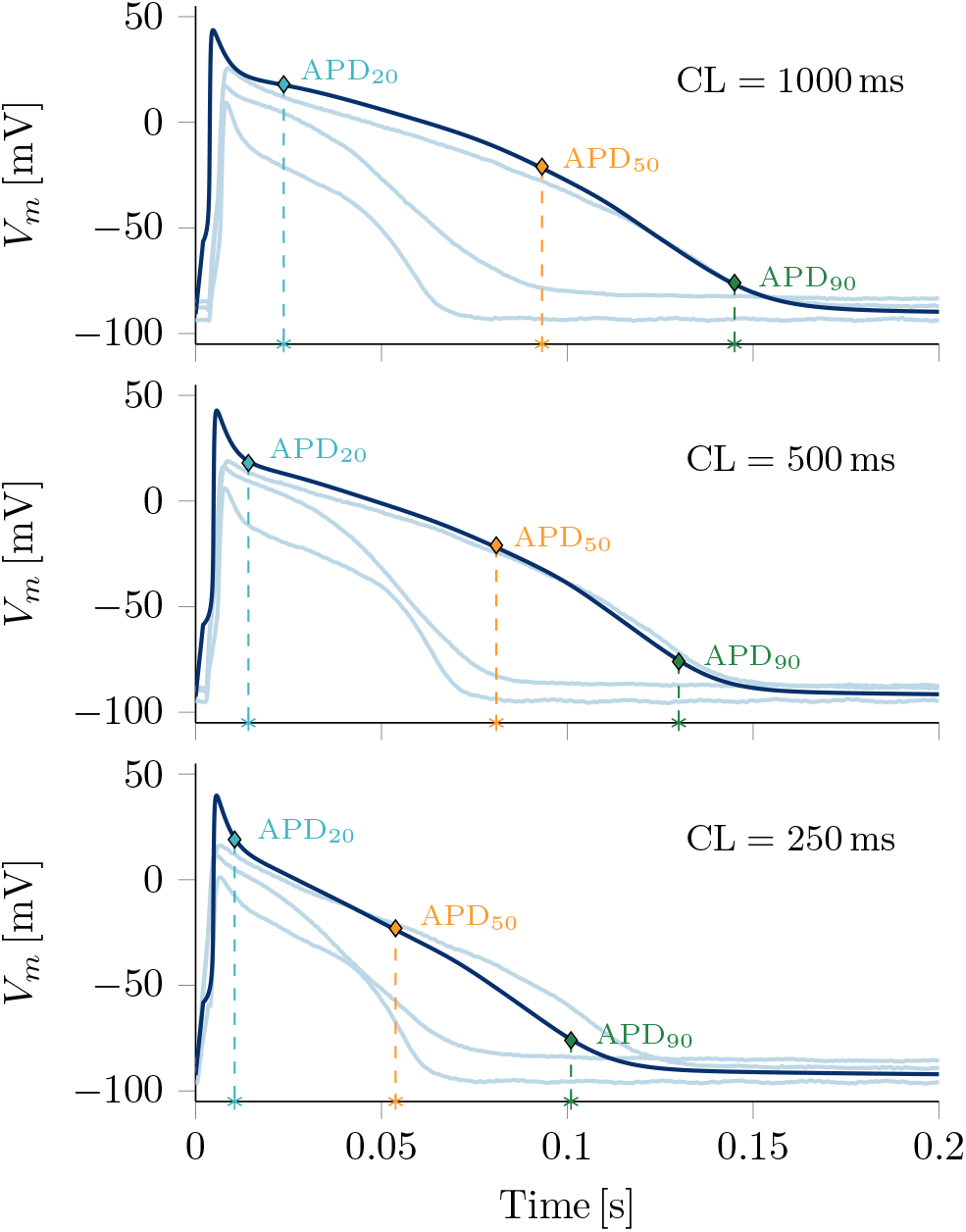
Experimental rate dependence traces. Illustrative experimental AP from three cells (light blue) and the AP simulated by the AL hiPSC model (dark blue) when considering three different frequencies. A qualitative analysis suggests that increased frequencies drive both the experimental and the simulated dynamic toward the reduction of the APD.

**Fig 6.**
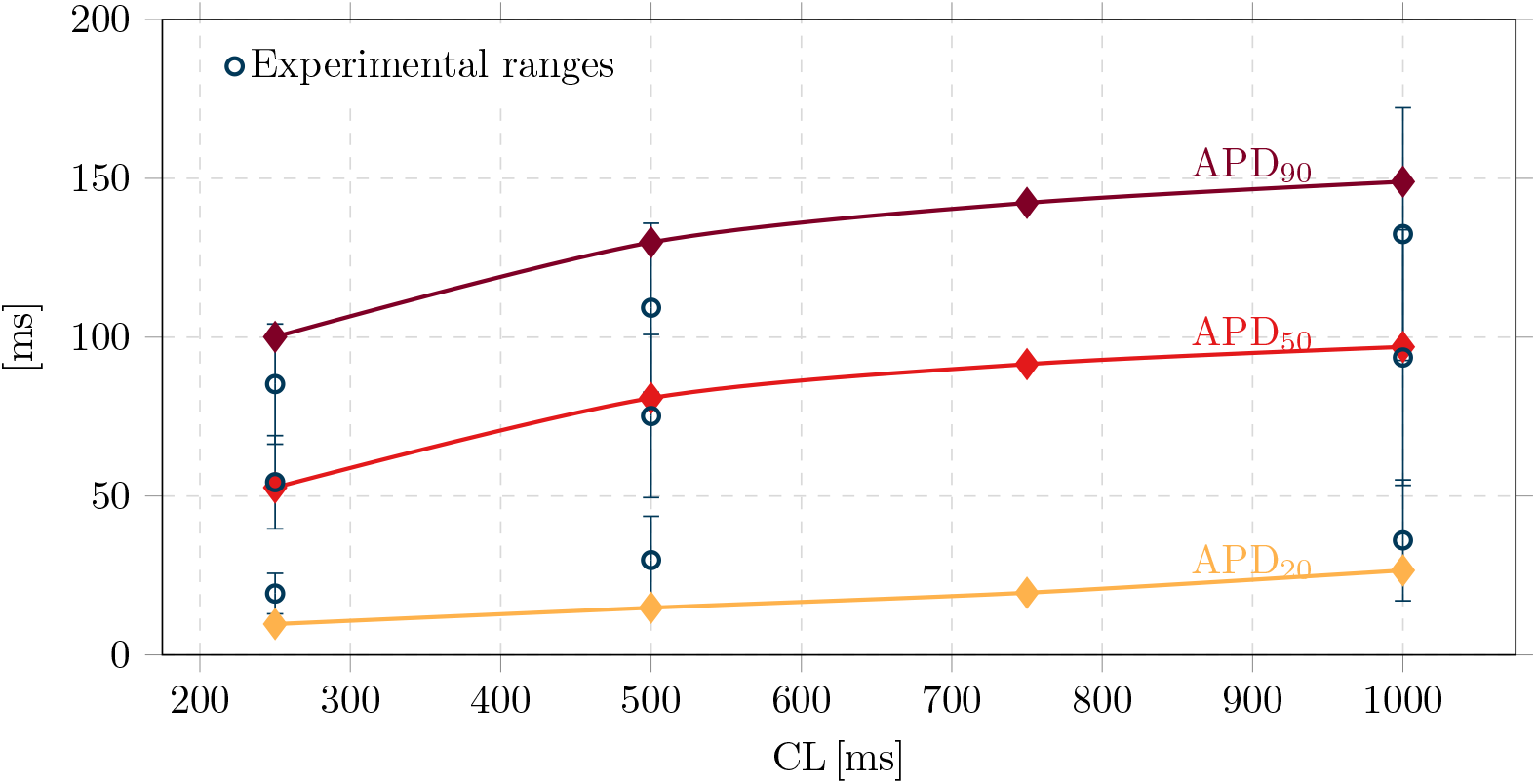
Rate dependence curves. Simulated APD_90_, APD_50_, and APD_20_ rate dependence curves (diamonds) produced by the novel AL model compared with *in–vitro* experimental data (circles and vertical error bars).

### 4–AP drug test

In order to compare the novel AL model with the experimental data, we challenged our model by exploiting the sensitivity of atrial *I*_*Kur*_ current to a selective blocker 4–aminopyridine (4–AP, 50 μM). In both isolated human adult atrial CMs and single atrial hiPSC–CM the blocking of the *I*_*Kur*_ resulted in APD_90_ prolongation [11]. As expected, in *in–vitro* experiments 4–AP superfusion causes the prolongation of AP in AL hiPSC–CMs. Analysis of 4–AP effects in APD changes (summarized in Table 6) showed that the highest prolongation was detected in the APD_20_ phase, where the *I*_*Kur*_ mostly contributed during the electrical activity of AL CMs.

**Table 6.** 4–AP effect on different AP phase. Experimental and simulated values of the biomarkers when considering 4–AP treatment. Experimental data provided as Mean*±*Std. Dev. in the form of Delta %, from [9].

| Biomarkers | Experimental delta % | Simulated delta % | Simulated absolute values |  |
| --- | --- | --- | --- | --- |
|  |  |  | Without 4AP | With 4AP |
| APD<sub>90</sub> [ms] | 19±21.187 | 17.19 | 148.94 | 174.5 |
| APD<sub>20</sub> [ms] | 56.8±63.87 | 36.2 | 26.60 | 36.2 |
| APD<sub>50</sub> [ms] | 36.4±37.63 | 20.09 | 96.94 | 116.41 |
| APD<sub>20/90</sub> | 0.3±0.31 | 15.9 | 0.179 | 0.207 |

We simulated *I*_*Kur*_ block by 50 μM of 4–AP as a 80% block of *I*_*Kur*_ maximum conductance, as suggested for human adult CMs in [20]. Similar values were published in [21] for human CMs, where dose–response curves show a 80% current block with 4–AP concentration of 50 μM. Human adult CMs are used as a reference since any information is provided about the *I*_*Kur*_ density in isolated hiPSC–CMs. Indeed, genes encoding for the *I*_*Kur*_ channel subunits are absent in the early phase of heart embryogenesis and are finely tuned in the developing heart, see [9]. Similarly, we expect that differentiating hiPSC–CM express mixed set of ion channels that affect the repolarization phase of AP in response to specific drug.

Our simulations suggest that *I*_*Kur*_ block induces an APD prolongation, in agreement with experimental results, as shown in Fig. 7. Furthermore, the quantitative comparison of APD on different AP phases in Table 6 validates the accuracy of the *in–silico* model, since every value APD change (delta %) is in the experimental variability range. Finally, also the higher contribution in the APD_20_ phase is preserved (36%).

**Fig 7.**
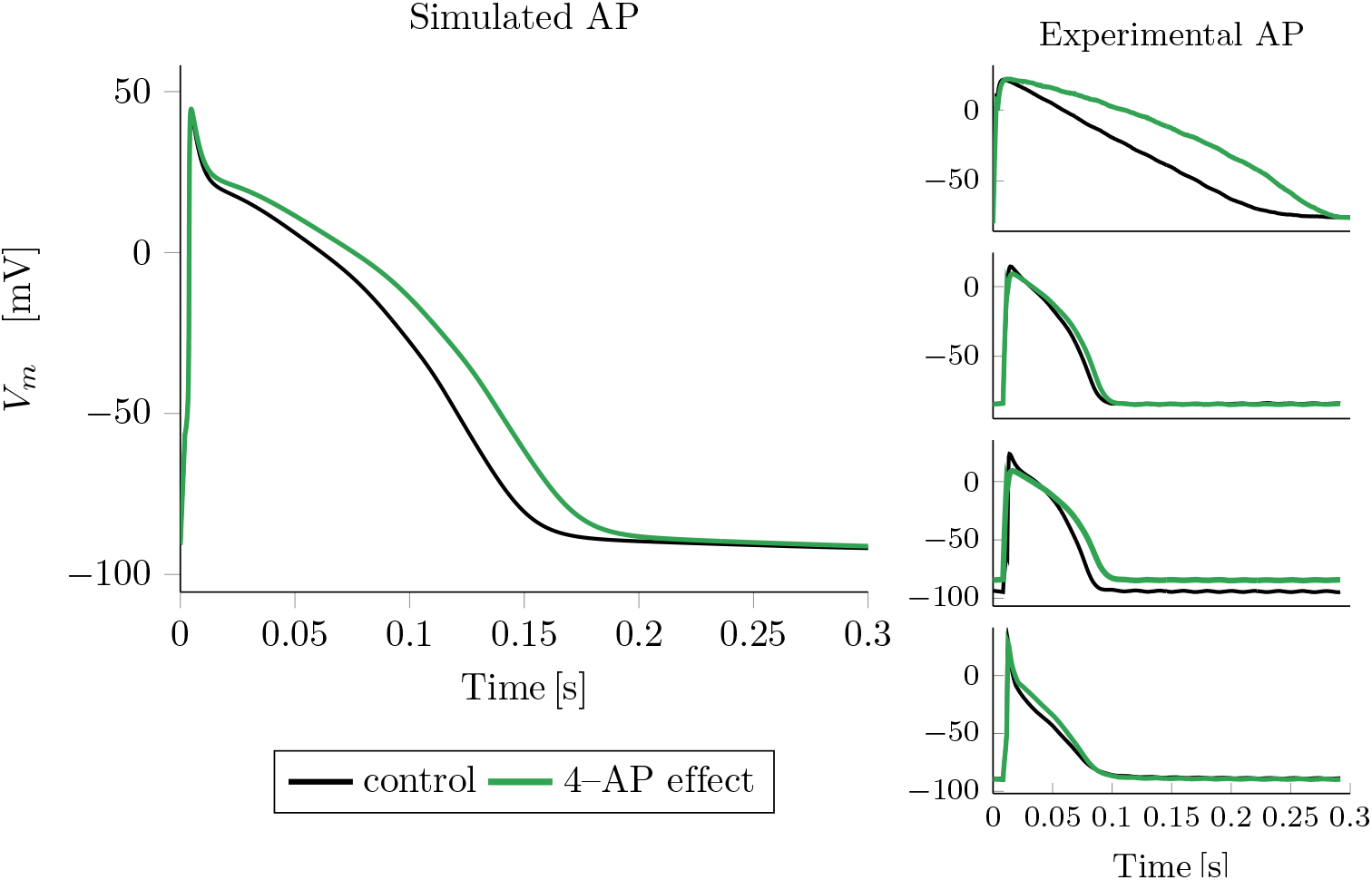
4–AP effect. Illustrative experimental AP from four cells (right side) and the AP simulated by the AL model (left side), when considering the control (black) or the 4–AP effect (green).

## Discussion

The focus on electrophysiological characteristics exhibited by AL cells allows a deeper understanding of the mechanisms underlying chamber–specific cardiomyopathies. Our work aimed to develop a highly specific AL hiPSC–CM model tailored to the distinct phenotype observed in atrial cells, thus contributing to elucidate the intricacies of atrial AP electrical abnormalities and acknowledge the broader implications for personalized medicine. This innovative approach leads us to recognize the pivotal role of modeling in advancing personalized drug testing methodologies, see e.g. [22].

Starting from an improved version of the ventricular Paci2020 hiPSC–CM model, [8] we developed and validated and AL model, which better recapitulates the phenotypical specificity. Two different VL pedigrees were published recently and both of them provided insights and contributions for the last Paci2020 model: Koivumäki2018 [23] and Kernik2019 [24], which were also based on the original Paci2013 hiPSC–CMs model. As suggested in [25], Koivumäki formulation employs a complex layered compartmental structure, which increases the computational cost of model simulations. Conversely, Kernik automaticity is sustained by a different *I*_*CaL*_ formulation, rather than being directly sustained by the Ca^2+^ handling dynamic. For these reasons, we started developing the new AL hiPSC–CMs model using Paci2020 as a parent model. Finally, this study also shows the predictive power of the model through an *in–silico* trial on an atrial–specific drug, in agreement with *in–vitro* data sets.

Research on hiPSC–CMs is rapidly developing, with new experimental data becoming available, which in turn serve as a driving force for the constantly evolving of computational models to offer more accurate *in–silico* tools. As for adult cardiac cells, which show different AP shapes according to their location and specialization in the heart, hiPSC-CMs show different AP morphologies, that are usually categorized as VL, AL, or nodal-like. In this paper we present an updated version of the ventricular Paci2020 model for AL hiPSC-CM, developed using literature data to identify the ionic currents most likely contributing to differentiate the VL from the AL AP in adult CMs. Because of the clinical interest in AF, we considered various K^+^ channels usually remodeled during AF simulations. Several of them are almost only expressed in the atria and they are summarized in [26]. Among them, we incorporated the atrial–specific *I*_*Kur*_ and *I*_*KCa*_ currents. Recent studies, such as [27, 28], suggest that native *I*_*Kur*_ density in AL hiPSC–CMs is substantially smaller than the *I*_*Kur*_ density of freshly isolated human atrial myocytes and a DC approach for the injection of virtual *I*_*Kur*_ current could lead the cell to a more adult phenotype. Future works will investigate *in–silico* the impact of an additional *I*_*Kur*_ current, also taking into account different formulations, such as Maleckar et al. [29]. Conversely, we did not take into account the Acetylcholine-sensitive K^+^ current (*I*_*K,ACh*_) and the Two-pore-domain K^+^ (*I*_*K*2*P*_) current. The first one is an atrial specific current activated by the neurotransmitter acetylcholine after the binding to the specific muscarinic receptor. For this reason its contribution to the AP in *in–vitro* isolated hiPSC-CMs can be neglected. The latter one is positively modulated in Schmidt’s model, [30], in the paroxysmal AF model and overexpressed in the chronic model. However in basal conditions the current does not concur in the electrical activity of a healthy patient with sinus rhythm. Furthermore, we choose to change the *I*_*K*1_ formulation, inspired by the Koivumäki 2014 model of human adult atrial CM. This latter change was implemented mainly to correctly take into account the use of DC technique in the experimental setup based on the injection of the Koivumäki *I*_*K*1_ current to electronically induce the maturation of *in–vitro* hiPSC–CMs, [9].

In order to keep the new model more consistent with experimental data and calibrate different parameters we have performed an optimization procedure, tightly bounded by the AP features. For the optimization of the model, we have built our cost function with the same biomarkers and experimental ranges provided by Altomare et al. [9], and we tuned parameters listed in Table 3. The result is a new AL hiPSC model where APs’ features in paced conditions match the Altomare et al. experimental dataset, as depicted in Fig. 4. The comparison between the experimental and simulated AP features reported in Table 4 shows how the model is matching the real APs. In particular, the AL model is fully compliant with the discriminating rule for the AL phenotype (APD_20*/*90_ 0.44). The only relevant discrepancy can be found in APA, with percentages of discrepancy equal to 10%. The mismatching in APA can be attributable to the high sensitivity of this biomarker on the external applied stimulus.

Our model was validated against the available experimental data ranging over three different pacing frequencies. The *in–vitro* experimental data set was not used in the calibration process, but the novel AL hiPSC model perfectly replicates the rate dependence curve, if considering the APD_50_ and APD_90_, as depicted in Fig. 6. The comparison between the provided and resulting AP features reported in Table 5, qualitatively overlapped in Fig. 5, shows how the model is matching the real experimental range. Also APD_20_ rate dependence curve qualitatively overlap experimental data, even if an accurate quantitative comparison (Table 5) highlights some negligible discrepancies for CL = 500 ms and CL = 250 ms.

We also validated our model when considering the response to the *I*_*Kur*_ atrial–specific current blocker 4–AP under an external stimulus; thus, simulation results were validated against the corresponding *in–vitro* experiments, which were not used during the calibration process (see Table 6). The rational we want to show is the key role of *I*_*Kur*_ current if considering AL hiPSC–CMs (and AL hiPSC models as well). The effect of current blocker on the AP shape and the resulting APD prolongation proving the phenothype specificity of in *in–vitro* cells, is perfectly replicated by the novel AL hiPSC *in–silico* model. In future works, we aim to test the rate dependence of the effects of *I*_*Kur*_ on AP repolarization in hiPSC–CMs.

As with all experimental models, the novel AL hiPSC is not without limitations. One limitation of our work could be the uniqueness of the data source: in fact we focused principally on the work of Altomare et al., [9], since it was the most comprehensive report about AL hiPSC–CMs maturation, classification and AP response to drugs. It must be noted that extensive experimental data sets from healthy mature hiPSC–CMs are nonexistent because different technologies for generating mature AP waveform are still under evaluation. We considered DC as a consolidated technique (see [28, 31, 32]), but new experimental data should be provided to improve the calibration of the model. Optimization approaches are recently being developed to improve the maturity of *in–vitro* hiPSC-CMs and bring them closer to an adult phenotype. Nowadays, the challenge of hiPSC–CM maturation has been tackled *in–silico*, trying to provide more adult virtual models. In this direction, our novel AL hiPSC model provides a useful tool to quantitatively predict the impact of phenotype specific drugs. Future perspectives absolutely consider the comparison with human adult models of atrial CMs and main differences with these models could suggest an operative direction towards the maturation of AL hiPSCs. Finally, an improved version of this AL hiPSC–CM model could take into account also the neglected atrial–specific currents *I*_*K,ACh*_ and *I*_*K*2*P*_, with the reasonable goal of simulating an engineered tissue in pathological and non basal conditions.

To sum up, in this work we present an updated and more specific version of an AL hiPSC-CM *in–silico* model, based on a new dataset of electrophysiological data and novel technologies to improve the cell maturation. Due to its relatively light formulation (24 ODEs), our model is suitable also for very large studies on *in–silico* populations, and future works will explore the possibility to support screening of different phenothype specific drug at various concentrations.

## Conclusion

In conclusion, this study introduces a novel set of phenotype-specific membrane currents and presents an *in–silico* model constrained by *in–vitro* experimental data to simulate the paced AP of matured AL hiPSC-CMs. Moreover, this model effectively recapitulates the electronic maturation process, transitioning from an unstable depolarized MDP to a hyperpolarized resting potential, and exhibits spontaneous firing activity in unpaced conditions. Finally, our model simulation accurately reflects the experimental rate dependence data and demonstrates the expected response to a specific current blocker.

## Supporting information

**S1 Appendix. Mathematical model**. This file contains the full set of equations of out novel AL hiPSC model and the list of constants and initial conditions at steady state. We highlighted in blue only the final changes in the Paci2020 model.

**S2 Appendix. List of currents**. This file contains the full list of membrane currents, exchanger and pumps, as well as the list of fluxes from the SR.

