## Appendix_1 for "A novel ionic model for matured and paced atrial–like hiPSC–CMs integrating *I*_*Kur*_ and *I*_*KCa*_ currents"

Appendix 1 – list of the equations

$V$: membrane potential in Volt.
$VmV$: membrane potential in milliVolt.
$t$: time simulation in seconds.

### 1 Constants

#### Extracellular ionic concentrations

$Na_o = 154.0$ (mM)
$K_o = 4.0$ (mM)
$Ca_o = 2.0$ (mM)

#### Cell size and dimensions

$C_m = 78.6672 \cdot 10^{-12}$ (F)
$V_c = 7012 \cdot 10^{-18}$ ($m^3$)
$V_{SR} = 465.199 \cdot 10^{-18}$ ($m^3$)

#### Maximum conductances and currents

$g_{Na} = 9.8001 \cdot 10^3$ (S/F)
$g_{NaL} = 13.3509$ (S/F)
$g_{CaL} = 7.45 \cdot 10^{-5}$ ($m^3/(F \times s)$)
$g_{to} = 59.45$ (S/F)
$g_{Kr} = 16.8723$ (S/F)
$g_{Ks} = 2.0856$ (S/F)
$g_{K1} = 0.169$ (S/F)
$g_f = 25.9$ (A/F)
$P_{NaK} = 2.2718$ (A/F)
$K_{NaCa} = 3450.7$ (A/F)
$I_{rel,max} = 75.4190$ (mM/s)
$V_{max,up} = 0.3226$ (mM/s)
$I_{leak,max} = 4.48209 \cdot 10^{-4}$ (1/s)
$g_{pCa} = 0.4570$ (A/F)
$g_{bNa} = 1.14$ (A/F)
$g_{bCa} = 0.114$ (S/F)
$\mathrm{coeff}_{\mathrm{Kur}} = 3.5$ (S/F)
$g_{KCa} = 0.0754$ (S/F)

#### Other constants

$Buf_c = 0.25$ (mM)
$Buf_{sr} = 10$ (mM)
$K_{buf_c} = 0.001$ (mM)
$K_{buf_{sr}} = 0.3$ (mM)
$K_{up} = 4.40425 \cdot 10^{-4}$ (mM)

$K_{pCa} = 0.0005$ (mM)
$F = 96485.3415$ (C/mol)
$R = 8.314472$ (J/(mol × K))
$T = 310$ (K)
$L_0 = 0.025$ (dimensionless)
$Q = 2.3$ (dimensionless)
$P_{kna} = 0.03$ (dimensionless)
$K_{sat} = 0.1$ (dimensionless)
$K_{mCa} = 1.38$ (mM)
$K_{mNai} = 87.5$ (mM)
$KCa_{on} = 47.0 \cdot 10^6$ (dimensionless)
$KCa_{off} = 13.0$ (dimensionless)
$\alpha = 2.16659$ (dimensionless)
$\gamma = 0.35$ (dimensionless)
$K_{mNa} = 40$ (mM)
$K_{mK} = 1$ (mM)
$RyR_{a1} = 0.1034$ ($\mu$M)
$RyR_{a2} = 0.050001$ ($\mu$M)
$RyR_{a,half} = 0.02632$ ($\mu$M)
$RyR_{o,half} = 0.00944$ ($\mu$M)
$RyR_{c,half} = 0.00167$ ($\mu$M)

### Initial conditions of state variables

$h_0 = 0.75$ (dimensionless)
$j_0 = 0.75$ (dimensionless)
$m_0 = 0$ (dimensionless)
$mL_0 = 0$ (dimensionless)
$hL_0 = 0.75$ (dimensionless)
$d_0 = 0$ (dimensionless)
$fCa_0 = 1$ (dimensionless)
$f1_0 = 1$ (dimensionless)
$f2_0 = 1$ (dimensionless)
$r_0 = 0$ (dimensionless)
$q_0 = 1$ (dimensionless)
$Xr1_0 = 0$ (dimensionless)
$Xr2_0 = 1$ (dimensionless)
$Xs_0 = 0$ (dimensionless)
$Xf_0 = 0.1$ (dimensionless)
$RyRa_0 = 0.3$ (dimensionless)
$RyRo_0 = 0.9$ (dimensionless)
$RyRc_0 = 0.1$ (dimensionless)
$V_0 = -0.070$ (V)
$Na_i = 9.2$ (mM)
$Ca_i = 0.0002$ (mM)
$Ca_{SR} = 0.32$ (mM)

### 2 Model's equations

#### Membrane Potential

$$\frac{\mathrm{d}V}{\mathrm{d}t} = -I_{ion} = -I_{K1} - I_{to} - I_{Kr} - I_{Ks} - I_{CaL} - I_{NaK} - I_{Na} - I_{NaL}$$
$$-I_{NaCa} - I_{pCa} - I_{f} - I_{Kur} - I_{KCa} - I_{bNa} - I_{bCa} + I_{stim}$$

#### $Na^+$ current, $I_{Na}$

$$I_{Na} = g_{Na} \cdot m^3 \cdot h \cdot j \cdot (V - E_{Na})$$

##### $I_{Na}$, $h$ gate

$$h_{inf} = \frac{1}{\sqrt{1 + \exp\left(\frac{VmV + 66.5}{6.8}\right)}}$$
$$\tau_h = 0.00007 + \frac{0.034}{1 + \exp\left(\frac{VmV+41}{5.5}\right) + \exp\left(-\frac{VmV+41}{14}\right)} + \frac{0.0002}{1 + \exp\left(-\frac{VmV+79}{14}\right)}$$
$$\frac{\mathrm{d}h}{\mathrm{d}t} = \frac{h_{inf} - h}{\tau_h}$$

##### $I_{Na}$, $j$ gate

$$j_{inf} = h_{inf}$$
$$\tau_j = 0.007 + \frac{1.5}{1 + \exp\left(\frac{VmV+41}{5.5}\right) + \exp\left(-\frac{VmV+41}{14}\right)} + \frac{0.02}{1 + \exp\left(-\frac{VmV+79}{14}\right)}$$
$$\frac{\mathrm{d}j}{\mathrm{d}t} = \frac{j_{inf} - j}{\tau_j}$$

##### $I_{Na}$, $m$ gate

$$m_{inf} = \frac{1}{1 + \exp\left(\frac{39+V}{-11.2}\right)}$$
$$\tau_m = 0.00001 + 0.00013 \cdot \exp\left(-\frac{(VmV + 48)^2}{15^2}\right) + \frac{0.000045}{1 + \exp\left(\frac{VmV+42}{-5}\right)}$$
$$\frac{\mathrm{d}m}{\mathrm{d}t} = \frac{m_{inf} - m}{\tau_m}$$

#### Late $Na^+$current, $I_{NaL}$

$$I_{Na} = g_{NaL} \cdot mL^3 \cdot hL \cdot (V - E_{Na})$$

##### $I_{NaL}$, $hL$ gate

$$hL_{inf} = \frac{1}{\left(1 + \exp\left(\frac{VmV+87.61}{7.488}\right)\right)}$$
$$\tau_{hL} = 0.2 \quad \text{(s)}$$
$$\frac{\mathrm{d}hL}{\mathrm{d}t} = \frac{hL_{inf} - hL}{\tau_{hL}}$$

#### $I_{NaL}$, $mL$ gate

$$mL_{inf} = \frac{1}{1 + \exp\left(\frac{-VmV - 42.85}{5.264}\right)}$$

$$\alpha_{mL} = \frac{1}{1 + \exp\left(\frac{-60 - VmV}{5}\right)}$$

$$\beta_{mL} = \frac{0.1}{1 + \exp\left(\frac{(VmV + 35)}{5}\right)} + \frac{0.1}{1 + \exp\left(\frac{(VmV - 50)}{200}\right)}$$

$$\tau_{mL} = \alpha_{mL} \cdot \beta_{mL}$$

$$\frac{\mathrm{d}mL}{\mathrm{d}t} = 1000 \cdot \frac{mL_{inf} - mL}{\tau_{mL}}$$

### L-type $Ca^{2+}$ current, $I_{CaL}$

$$I_{CaL} = \frac{4 \cdot V \cdot F^2 \cdot g_{CaL}}{R \cdot T} \cdot \frac{\left(Ca_i \cdot \exp\left(\frac{2 \cdot V \cdot F}{R \cdot T}\right) - 0.341 \cdot Ca_o\right)}{\exp\left(\frac{2 \cdot V \cdot F}{R \cdot T}\right) - 1} \cdot d \cdot f1 \cdot f2 \cdot fCa$$

#### $I_{CaL}$, $d$ gate

$$d_{inf} = \frac{1}{1 + \exp\left(\frac{-VmV - 9.1}{7}\right)}$$

$$\alpha_d = \frac{1.4}{1 + \exp\left(\frac{-35 - VmV}{13}\right)} + 0.25$$

$$\beta_d = \frac{1.4}{1 + \exp\left(\frac{VmV + 5}{5}\right)}$$

$$\gamma_d = \frac{1}{1 + \exp\left(\frac{50 - VmV}{20}\right)}$$

$$\tau_d = \alpha_d \cdot \beta_d + \gamma_d$$

$$\frac{\mathrm{d}d}{\mathrm{d}t} = 1000 \cdot \frac{d_{inf} - d}{\tau_d}$$

#### $I_{CaL}$, $f_{Ca}$ gate

$$\alpha_{fCa} = \frac{1}{1 + \left(\frac{Ca_i}{0.0006}\right)^8}$$

$$\beta_{fCa} = \frac{0.1}{1 + \exp\left(\frac{Ca_i - 0.0009}{0.0001}\right)}$$

$$\gamma_{fCa} = \frac{0.3}{1 + \exp\left(\frac{Ca_i - 0.00075}{0.0008}\right)}$$

$$fCa_{inf} = \frac{\alpha_{fCa} + \beta_{fCa} + \gamma_{fCa}}{1.3156}$$

$$\tau_{fCa} = 0.002 \quad \text{(s)}$$

$$\frac{\mathrm{d}fCa}{\mathrm{d}t} = \mathrm{const_{fCa}} \cdot \frac{fCa_{inf} - fCa}{\tau_{fCa}}$$

$$\mathrm{const_{fCa}} = \begin{cases} 0, & \text{if} \quad fCa_{inf} > fCa \text{ and } VmV > -60 \quad \text{(mV)} \\ 1, & \text{otherwise} \end{cases}$$

#### $I_{CaL}$, *f1* gate

$$f1_{\text{inf}} = \frac{1}{1 + \exp\left(\frac{VmV+26}{3}\right)}$$

$$\tau_{f1} = \left(20 + 1102.5 \cdot \exp\left(-\frac{(V+50)^2}{15}\right) + \frac{200}{1+\exp\left(\frac{13-V}{10}\right)} + \frac{280}{1+\exp\left(\frac{30+V}{10}\right)}\right) \cdot \text{const}_{\text{f1}}$$

$$\text{const}_{\text{f1}} = \begin{cases} 1.35 \cdot \left[1 + 1433 \cdot \left(Ca_i - 50 \cdot 10^{-6}\right)\right], & \text{if } \frac{df1}{dt} > 0 \\ 1, & \text{otherwise} \end{cases}$$

$$\frac{\mathrm{d}f1}{\mathrm{d}t} = 1000 \cdot \frac{f1_{inf} - f1}{\tau_{f1}}$$

#### $I_{CaL}$, *f2* gate

$$f2_{inf} = \frac{0.67}{1 + \exp\left(\frac{VmV+32}{4}\right)} + 0.33$$

$$\tau_{f2} = 600 \cdot \exp\left(-\frac{(V+50)^2}{400}\right) + \frac{31}{1+\exp\left(\frac{25-V}{10}\right)} + \frac{1}{1+\exp\left(\frac{30+V}{10}\right)}$$

$$\frac{\mathrm{d}f2}{\mathrm{d}t} = 1000 \cdot \frac{f2_{inf} - f2}{\tau_{f2}}$$

### Transient outward current, $I_{to}$

$$I_{to} = g_{to} \cdot r \cdot q \cdot (V - E_K)$$

#### $I_{to}$, *r* gate

$$r_{inf} = \frac{1}{1 + \exp\left(\frac{22.3 - VmV}{18.75}\right)}$$

$$\tau_r = \frac{14.40516}{1.037 \cdot e^{0.09 \cdot (VmV + 30.61)} + 0.369 \cdot e^{-0.12 \cdot (VmV + 23.84)}} + 2.75352$$

$$\frac{\mathrm{d}r}{\mathrm{d}t} = 1000 \cdot \frac{r_{inf} - r}{\tau_r}$$

#### $I_{to}$, *q* gate

$$q_{inf} = \frac{1}{1 + \exp\left(\frac{VmV+53}{13}\right)}$$

$$\tau_q = \frac{39.102}{0.57 \cdot \exp\left(-0.08 \cdot (VmV + 44)\right) + 0.065 \cdot \exp\left(0.1 \cdot (VmV + 45.93)\right)} + 6.06$$

$$\frac{\mathrm{d}q}{\mathrm{d}t} = 1000 \cdot \frac{q_{inf} - q}{\tau_q}$$

### Rapid delayed rectifier $K^+$ current, $I_{Kr}$

$$I_{Kr} = g_{Kr} \cdot \sqrt{\frac{K_o}{5.4}} \cdot Xr1 \cdot Xr2 \cdot (V - E_K)$$

#### $I_{Kr}$, *Xr1* gate

$$V_{half,Xr1} = 1000 \cdot \left( -\frac{R \cdot T}{F \cdot Q} \cdot \ln \left( \frac{\left(1 + \frac{Ca_o}{2.6}\right)^4}{L_0 \cdot \left(1 + \frac{Ca_o}{0.58}\right)^4} \right) - 0.019 \right)$$

$$Xr1_{inf} = \frac{1}{1 + \exp\left(\frac{V_{half,Xr1} - VmV}{4.9}\right)}$$

$$\alpha_{Xr1} = \frac{450}{1 + \exp\left(\frac{-45 - VmV}{10}\right)}$$

$$\beta_{Xr1} = \frac{6}{1 + \exp\left(\frac{VmV + 30}{11.5}\right)}$$

$$\tau_{Xr1} = \alpha_{Xr1} \cdot \beta_{Xr1}$$

$$\frac{\mathrm{d}Xr1}{\mathrm{d}t} = 1000 \cdot \frac{Xr1_{inf} - Xr1}{\tau_{Xr1}}$$

#### $I_{Kr}$, *Xr2* gate

$$Xr2_{inf} = \frac{1}{1 + \exp\left(\frac{VmV + 88}{50}\right)}$$

$$\alpha_{Xr2} = \frac{3}{1 + \exp\left(\frac{-60 - VmV}{20}\right)}$$

$$\beta_{Xr2} = \frac{1.12}{1 + \exp\left(\frac{VmV - 60}{20}\right)}$$

$$\tau_{Xr2} = \alpha_{Xr2} \cdot \beta_{Xr2}$$

$$\frac{\mathrm{d}Xr2}{\mathrm{d}t} = 1000 \cdot \frac{Xr2_{inf} - Xr2}{\tau_{Xr2}}$$

### Slow delayed rectifier $K^+$ current, $I_{Ks}$

$$I_{Ks} = g_{Ks} \cdot Xs^2 \cdot \left( 1 + \frac{0.6}{1 + \left(\frac{3.8 \cdot 10^{-5}}{Ca_i}\right)^{1.4}} \right) \cdot (V - E_{Ks})$$

#### $I_{Ks}$, *Xs* gate

$$Xs_{inf} = \frac{1}{1 + \exp\left(\frac{-20 - VmV}{16}\right)}$$

$$\alpha_{Xs} = \frac{1100}{\sqrt{1 + \exp\left(\frac{-10 - VmV}{6}\right)}}$$

$$\beta_{Xs} = \frac{1}{1 + \exp\left(\frac{VmV - 60}{20}\right)}$$

$$\tau_{Xs} = \alpha_{Xs} \cdot \beta_{Xs}$$

$$\frac{\mathrm{d}Xs}{\mathrm{d}t} = 1000 \cdot \frac{Xs_{inf} - Xs}{\tau_{Xs}}$$

### Inward rectifier $K^+$ current, $I_{K1}$

$$I_{K1} = g_{K1} \cdot K_i^{0.4457} \cdot \frac{V - E_K}{1.0 + e^{1.5(V - E_K + 3.6)F/RT}}$$

### Hyperpolarization activated funny current, $I_f$

$$f_{Na} = 0.37$$
$$f_{Na} = 1 - f_{Na}$$
$$I_{fK} = f_K \cdot G_f \cdot (V - E_K)$$
$$I_{fNa} = f_{Na} \cdot G_f \cdot Xf_{inf} \cdot (V - E_{Na})$$
$$I_f = I_{fK} + I_{fNa}$$

#### $I_f$, $Xf$gate

$$Xf_{inf} = \frac{1}{1 + \exp\left(\frac{VmV+69}{8}\right)}$$
$$\tau_{Xf} = \frac{5600}{1 + \exp\left(\frac{VmV+65}{7}\right)} + \exp\left(\frac{VmV + 65}{19}\right)$$
$$\frac{\mathrm{d}Xf}{\mathrm{d}t} = 1000 \cdot \frac{Xf_{inf} - Xf}{\tau_{Xf}}$$

### $\mathrm{Na}^+$ component of $I_f$

$$I_{fNa} = 0.42 \cdot g_f \cdot Xf \cdot (V - E_{Na})$$

### $\mathrm{Na}^+/\mathrm{K}^+$ pump current, $I_{NaK}$

$$I_{NaK} = P_{NaK} \cdot \frac{\frac{K_o}{K_o+K_{mK}} \cdot \frac{Na_i}{Na_i+K_{mNa}}}{1 + 0.1245 \cdot \exp\left(\frac{-0.1 \cdot V \cdot F}{R \cdot T}\right) + 0.0353 \cdot \exp\left(\frac{-V \cdot F}{R \cdot T}\right)}$$

### $\mathrm{Na}^+/\mathrm{Ca}^{2+}$ exchanger current, $I_{NaCa}$

$$I_{\mathrm{NaCa}} = \frac{K_{\mathrm{NaCa}} \cdot \left(\exp\left(\frac{\gamma \cdot V \cdot F}{R \cdot T}\right) \cdot Na_i^3 \cdot \mathrm{Ca}_o - \exp\left(\frac{(\gamma-1) \cdot V \cdot F}{R \cdot T}\right) \cdot Na_o^3 \cdot Ca_i \cdot \alpha\right)}{(K_{mNai}^3 + Na_o^3) \cdot (K_{mCa} + Ca_o) \cdot \left(1 + K_{sat} \cdot \exp\left(\frac{(\gamma-1)VF}{R \cdot T}\right)\right)}$$

### Small conductance $\mathrm{Ca}^{2scii+}$ activated $\mathrm{K}^+$ channel, $I_{KCa}$

$$\frac{\mathrm{d}O}{\mathrm{d}t} = (1 - O) \cdot KCa_{on} \cdot Ca_i^2 - O \cdot KCa_{off}$$
$$I_{KCa} = g_{KCa} \cdot O \cdot \frac{1}{1 + \exp(\frac{V - E_K \cdot 10^3 + 120.0}{45.0})}(V - E_K \cdot 10^3)$$

### Ultrarapid delayed rectifier current, $I_{Kur}$

$$I_{Kur} = g_{Kur} \cdot u_a^3 \cdot u_i \cdot \left(V - E_K \cdot 10^3\right)$$

$$g_{Kur} = \text{coeff}_{\text{Kur}} \cdot \left[0.005 + \frac{0.05}{1.0 + \exp\left(-\frac{V-15.0}{13.0}\right)}\right]$$

$$\alpha_{u(a)} = 0.65 \left[\exp\left(-\frac{V+10.0}{8.5}\right) + \exp\left(-\frac{V-30.0}{59.0}\right)\right]^{-1}$$

$$\beta_{u(a)} = 0.65 \cdot \left[2.5 + e^{-\frac{V+82.0}{17.0}}\right]^{-1}$$

$$\tau_{u(a)} = \frac{K_{Q,10}}{\alpha_{u(a)} + \beta_{u(a)}}$$

$$u_{a(\infty)} = \left[1.0 + \exp\left(-\frac{V+30.0}{9.6}\right)\right]^{-1}$$

$$\frac{du_a}{dt} = \frac{u_{a(\infty)} - u_a}{\tau_{u(a)}}$$

$$\alpha_{u(i)} = \left[21.0 + \exp\left(-\frac{V-185.0}{28.0}\right)\right]^{-1}$$

$$\beta_{u(i)} = \exp\left(-\frac{V+158.0.0}{16.0}\right)$$

$$\tau_{iur} = \frac{K_{Q,10}}{\alpha_{u(i)} + \beta_{u(i)}}$$

$$u_{i(\infty)} = \left[1.0 + \exp\left(-\frac{V-99.45}{27.48}\right)\right]^{-1}$$

$$\frac{du_i}{dt} = \frac{u_{i(\infty)} - u_i}{\tau_{u(i)}}$$

### $Ca^{2+}$ pump current, $I_{pCa}$

$$I_{pCa} = \frac{g_{pCa} \cdot Ca_i}{Ca_i + K_{pCa}}$$

### $Na^{+}$ dynamics

$$\frac{\mathrm{d}Na_i}{\mathrm{dt}} = -C_m \cdot \frac{I_{Na} + I_{NaL} + I_{fNa} + I_{bNa} + 3I_{NaK} + 3I_{NaCa}}{F \cdot V_c}$$

### Background currents

$$I_{bNa} = g_{bNa} \cdot (V - E_{Na})$$
$$I_{bCa} = g_{bCa} \cdot (V - E_{Ca})$$

### $\mathbf{Ca^{2+}}$ dynamics

$$RyR_{CaSR} = 1 - \frac{1}{1 + \exp\left(\frac{Ca_{SR} - 0.3}{0.1}\right)}$$

$$RyRa_{inf} = RyR_{a1} - \frac{RyR_{a2}}{1 + \exp\left(\frac{1000 \cdot Ca_i - RyR_{a,half}}{0.0082}\right)}$$

$$\tau_{RyRa} = 1\,(\mathrm{s})$$

$$\frac{\mathrm{d}RyRa}{\mathrm{d}t} = \frac{RyRa_{inf} - RyRa}{\tau_{RyRa}}$$

$$RyRo_{inf} = 1 - \frac{1}{1 + \exp\left(\frac{1000 \cdot Ca_i - (RyRa + RyR_{o,half})}{0.003}\right)}$$

$$\tau_{RyRo} = \begin{cases} 18.75, & \text{if } (RyRo_{inf} \geq RyRo) \\ 1.875, & \text{otherwise} \end{cases}$$

$$\frac{\mathrm{d}RyRo}{\mathrm{d}t} = 1000 \cdot \frac{RyRo_{inf} - RyRo}{\tau_{RyRo}}$$

$$RyRc_{inf} = \frac{1}{1 + \exp\left(\frac{1000 \cdot Ca_i - (RyRa + RyR_{c,half})}{0.001}\right)}$$

$$\tau_{RyRc} = \begin{cases} 175.0, & \text{if } (RyRc_{inf} \geq RyRc) \\ 87.5, & \text{otherwise} \end{cases}$$

$$\frac{\mathrm{d}RyRc}{\mathrm{d}t} = 1000 \cdot \frac{RyRc_{inf} - RyRc}{\tau_{RyRc}}$$

$$I_{rel} = I_{rel,max} \cdot RyR_{CaSR} \cdot RyRo \cdot RyRc \cdot (Ca_{SR} - Ca_i)$$

$$I_{up} = \frac{V_{max,up}}{1 + \frac{K_{up}^2}{Ca_i^2}}$$

$$I_{leak} = I_{leak,max} \cdot (Ca_{SR} - Ca_i)$$

$$Ca_{i_{bufc}} = \frac{1}{1 + \frac{Buf_c \cdot K_{buf_c}}{(Ca_i + K_{buf_c})^2}}$$

$$Ca_{sr_{bufsr}} = \frac{1}{1 + \frac{Buf_{sr} \cdot K_{buf_{sr}}}{(Ca_{SR} + K_{buf_{sr}})^2}}$$

$$\frac{\mathrm{d}Ca_i}{\mathrm{d}t} = Ca_{i_{bufc}} \cdot \left(I_{leak} - I_{up} + I_{rel} - \frac{(I_{CaL} + I_{bCa} + I_{p_{Ca}} - 2 \cdot I_{NaCa})}{2 \cdot V_c \cdot F} \cdot C_m\right)$$

$$\frac{\mathrm{d}Ca_{SR}}{\mathrm{d}t} = \frac{Ca_{sr_{bufs}} \cdot V_c}{V_{sr}} \cdot (I_{up} - (I_{rel} + I_{\mathrm{leak}}))$$

### Reversal potentials

$$E_{Na} = \frac{R \cdot T}{F} \cdot \ln \frac{Na_o}{Na_i}$$

$$E_K = \frac{R \cdot T}{F} \cdot \ln \frac{K_o}{K_i}$$

$$E_{Ks} = \frac{R \cdot T}{F} \cdot \ln \frac{K_o + P_{kna} \cdot Na_o}{K_i + P_{kna} \cdot Na_i}$$

$$E_{Ca} = \frac{0.5 \cdot R \cdot T}{F} \cdot \ln \frac{Ca_o}{Ca_i}$$

$$E_f = -0.017 \qquad (\mathrm{V})$$

### Stimulus current, $I_{stim}$

$I_{app} = 1.41 \cdot 10^{-9} \qquad \text{(A)}$

$I_{stim,period} = 1$

$I_{stim,duration} = 2 \qquad \text{(ms)}$

$$\mathrm{const_{f1}} = \begin{cases} I_{app}, & \text{if} \quad 0 \leq t - \lfloor \frac{t}{I_{stim,period}} \rfloor \leq I_{stim,duration} \\ 0, & \text{otherwise} \end{cases}$$
