## Appendix_2 for "A novel ionic model for matured and paced atrial–like hiPSC–CMs integrating *I*_*Kur*_ and *I*_*KCa*_ currents"

Appendix 2 – list of the currents

### 1 Membrane currents

$I_{Na}$ $Na^+$ current

$I_{NaL}$ late $Na^+$ current

$I_f$ funny current

$I_{CaL}$ L-type $Ca^{2+}$ current

$I_{to}$ transient outward $K^+$ current

$I_{Kr}$ rapid delayed rectifier $K^+$ currents

$I_{Ks}$ slow delayed rectifier $K^+$ currents

$I_{NaCa}$ $Na^+/Ca^{2+}$ exchanger

$I_{NaK}$ $Na^+/K^+$ pump

$I_{pCa}$ sarcolemmal $Ca^{2+}$ pump

$I_{bNa}$ $Na^+$ background current

$I_{bCa}$ $Ca^{2+}$ background current

$I_{K1}$ $Ca^{2+}$ time independent inward–rectifier $K^+$ current

$I_{Kur}$ $Ca^{2+}$ ultrarapid delayed rectifier current

$I_{KCa}$ $Ca^{2+}$ small conductance $Ca^{2+}$ activated $K^+$ channel current

### 2 Fluxes from the SR

$I_{rel}$ RyR–sensitive release current

$I_{up}$ Sarco–Endoplasmic Reticulum $Ca^{2+}$ ATPase (SERCA) pump

$I_{leak}$ leakage current
